# LytM family endopeptidase LmdC is controlled by regulated auto processing and proteolytic destruction in *Pseudomonas aeruginosa*

**DOI:** 10.64898/2026.10.09.758143

**Authors:** Alexis G. Sommerfield, Andrew J. Darwin

**Author notes:** Address correspondence to: Andrew J. Darwin.

## Abstract

Enzymes that cleave the bacterial peptidoglycan sacculus, known collectively as autolysins, are critical for growth, division and morphogenesis, but their activities must be carefully controlled to avoid catastrophic lysis. Some autolysin control mechanisms have emerged, but we still know little about how most of them are regulated. Here, we report a novel control mechanism for the LytM/M23 family endopeptidase PA4404 in *P. aeruginosa*, which we have named LmdC. We have discovered that LmdC is auto catalytically cleaved into N- and C-terminal domains that are then degraded by the carboxyl-terminal processing proteases Prc and CtpA respectively. The effects of various mutations on LmdC-linked β-lactam resistance suggested that full length LmdC is the functional enzyme, with its auto catalytic cleavage serving only to initiate its proteolytic destruction. The auto catalytic cleavage of LmdC was inhibited by overproducing a polymer-forming cytoskeletal protein bactofilin homolog encoded by PA0664, which we named Baf, and a protein of unknown function encoded by PA0663, which we named Api (<u>a</u>uto <u>p</u>rocessing inhibitor). We hypothesize that Baf and Api could promote a location specific role for LmdC. In this scenario, the dangerous potential for any free LmdC to cleave peptidoglycan at the wrong place would be prevented by its immediate auto processing and degradation. The widespread co-occurrence and linkage of *lmdC* and *baf* homologs, coupled with conserved features of those LmdC homologs, makes it likely that regulated LmdC auto processing is a widely conserved control mechanism.

**IMPORTANCE:** A peptidoglycan cell wall is an essential component of almost all bacterial cells and antibiotics that interfere with its synthesis have been critical infection treatments for decades. Peptidoglycan must also be cleaved to allow incorporation of new material for growth and division. However, the autolysin enzymes that cleave peptidoglycan must be controlled to prevent cell rupture. It is not known how this control is achieved in most cases, but it is an important area of research because these control mechanisms could be targets for therapeutic intervention. This study describes the discovery of regulated auto catalytic processing as an autolysin control mechanism in the human pathogen *Pseudomonas aeruginosa*, which is also likely to be widely conserved in other species.

## INTRODUCTION

*Pseudomonas aeruginosa* is a widespread Gram-negative bacterium that causes serious opportunistic human infections (1). It is a leading cause of health care setting-acquired disease, and The World Health Organization has declared it a “Priority 1: Critical” pathogen (2, 3). Increasing antibiotic resistance and outbreaks of multidrug resistant strains in hospitals underlines the need for more study and the identification of new therapeutic targets (1, 4). The cell envelope of almost all bacteria contains a mesh-like cell wall (sacculus) composed of peptidoglycan. Peptidoglycan consists of linear glycan strands of alternating N-acetyl muramic acid and N-acetyl glucosamine disaccharide units, with a short amino acid peptide chain attached to N-acetyl muramic acid (5, 6). Covalent bonds between peptide chains of adjacent glycan strands result in an extensively cross-linked cell wall that withstands turgor pressure. Drugs that inhibit the formation of these cross-links have been a mainstay in the clinic for decades. New peptidoglycan must be synthesized for cell elongation and division, but the existing peptidoglycan mesh must also be cleaved to incorporate the new material (7, 8). Peptidoglycan cleavage is also important for morphogenesis and for the assembly of trans-envelope structures such as secretion systems and flagella (9–11).

Various enzymes, collectively called autolysins, can cleave the different bonds within intact peptidoglycan, with other enzymes acting on some of the resulting soluble products (9–11). Autolysin activities must be carefully controlled to avoid lethal rupture of the sacculus. This makes research into autolysin control mechanisms important, with the potential to identify new therapeutic targets. For most autolysins, there is either no information, or incomplete information about how they are regulated. However, while there are many more questions than answers, some control mechanisms have begun to emerge (12). These include modification of the peptidoglycan substrate, trans-acting regulators, and the control of autolysin concentration by various mechanisms, including degradation by carboxyl-terminal processing proteases (CTPs) (13, 14).

An *Escherichia coli* CTP named Prc forms a complex with the outer membrane lipoprotein NlpI to degrade several autolysins (13, 15–18). We found a remarkable parallel between Prc in *E. coli* and a CTP named CtpA in *P. aeruginosa* (19). Like *E. coli* Prc, CtpA forms a complex with an outer membrane lipoprotein (LbcA) and this LbcA•CtpA complex degrades four PG cross-link endopeptidases (14, 20). In both species, the lipoprotein binding partner acts as a scaffold that binds to the CTP and its substrates independently, and the proteolytic activity of the complex is regulated (19, 21–24). However, CtpA and Prc are in divergent CTP subgroups, their lipoprotein binding partners are not homologous, and the complexes have major structural differences (17, 25, 26). The functional parallel between *P. aeruginosa* CtpA and *E. coli* Prc is also surprising because in addition to CtpA, *P. aeruginosa* has a homolog of *E. coli* Prc. However, the only described role for *P. aeruginosa* Prc is to degrade a mutant form of anti-sigma factor MucA, triggering the mucoid conversion phenotype that occurs in individuals with cystic fibrosis (27–30).

We have become interested to discover the endogenous roles of Prc in *P. aeruginosa*. In this study, we report that one of the CtpA substrates is also degraded by Prc. Our investigations into this phenomenon have uncovered a novel autolysin control mechanism. The PA4404 endopeptidase, which we have named LmdC, is auto catalytically cleaved into N- and C-terminal domains that are degraded by Prc and CtpA respectively. The auto catalytic cleavage of LmdC appears to be controlled by two proteins that co-occur with LmdC homologs in some *Pseudomonas* species. One is the cytoskeletal protein bactofilin and the other is an integral inner membrane protein of previously unknown function. This autolysin control mechanism is likely to be conserved in many of the LmdC homologs that occur widely in diverse bacterial species.

## RESULTS

### N- and C-terminal fragments of PA4404 (LmdC) accumulate differentially in *prc* and *ctpA* null mutants

At the start of this study, the only demonstrated substrate of Prc in *P. aeruginosa* was the mutant MucA22 protein, which is often selected during lung infections in individuals with cystic fibrosis (30). The wild type MucA protein is not a Prc substrate, so to begin to search for native Prc substrates, we first investigated if Prc affected any of the known CtpA substrates (14, 20). We used our polyclonal antisera to compare the effect of *prc* and *ctpA* null mutations on steady state levels and found that one of the CtpA substrates, PA4404, had an intriguing accumulation pattern. PA4404 is the only member of the LmdC (<u>L</u>yt<u>M</u> <u>d</u>omain protein <u>C</u>) clade of lysostaphin-type LytM/MEROPS M23 endopeptidases in *P. aeruginosa* and so we will refer to it as LmdC (31). A band corresponding to the predicted 33 kDa molecular weight of LmdC was detected, but it did not appear to accumulate in the Δ*prc* or Δ*ctpA* mutants (Fig. 1a). However, an ∼12 kDa LmdC protein accumulated in a Δ*prc* strain, an ∼22 kDa LmdC protein accumulated in a Δ*ctpA* strain, and both accumulated in a Δ*prc* Δ*ctpA* strain (Fig. 1a).

**FIG 1.**
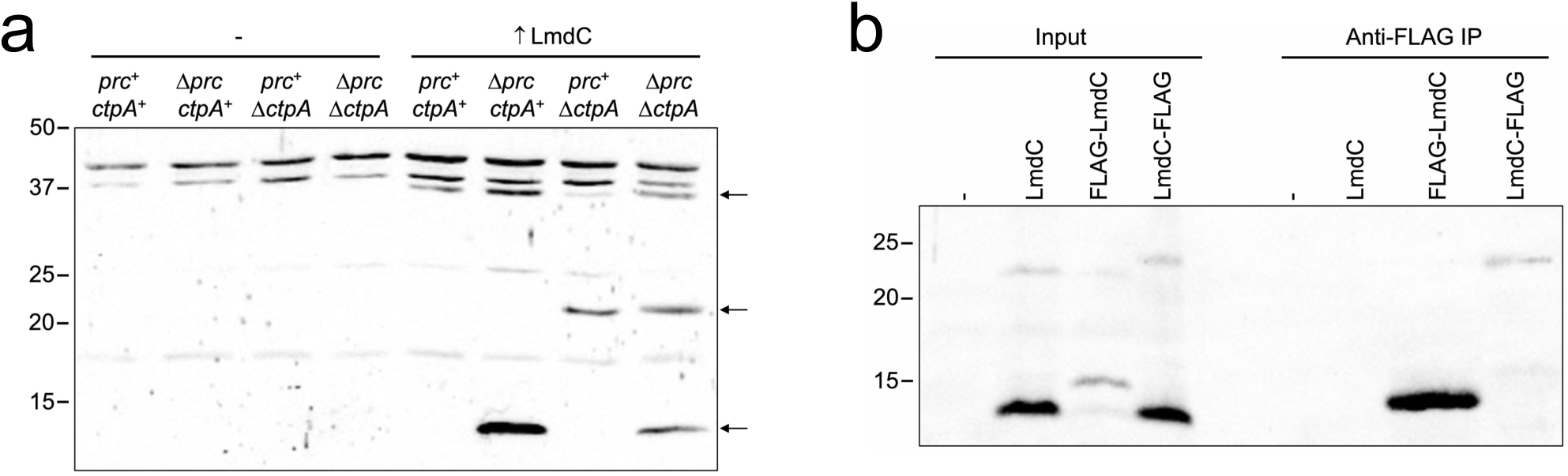
Fragments of LmdC accumulate differentially in Δ*prc* and Δ*ctpA* mutants. (a) Anti-LmdC immunoblot of whole-cell lysates of Δ*lmdC* strains containing a plasmid encoding LmdC (↑LmdC) or the empty vector (-). Arrows on the right indicate LmdC-specific protein bands, whereas all other bands are cross-reactive with the polyclonal antiserum (i.e., present in the empty vector strains). (b) Evidence that the LmdC fragments are N- and C-terminal domains. Anti-LmdC immunoblot of input lysates and immunoprecipitates (Anti-FLAG IP) from Δ*lmdC* strains containing a plasmid encoding LmdC, FLAG-LmdC, LmdC-FLAG or the empty vector (-). For both panels approximate sizes of molecular-mass marker proteins (M), in kilodaltons, are on the left. Data presented are single representatives of replicate experiments.

We noticed previously that a 20-25 kDa LmdC-FLAG protein accumulated in a Δ*ctpA* strain when detected by anti-FLAG immunoblot, even though the predicted molecular weight of LmdC is ∼33 kDa (14). We did not understand the reason for the size discrepancy at that time, but we now realized that the ∼12 and 22 kDa proteins accumulating in Δ*prc* and Δ*ctpA* mutants could together account for the expected mass of LmdC. Our previous anti-FLAG detection of the 20-25 kDa LmdC-FLAG protein suggested that the ∼ 22 kDa protein had the C-terminus (14). Therefore, we hypothesized that the ∼ 12 kDa protein had the N-terminus. To test this, we did anti-FLAG immunoprecipitations from a Δ*prc* Δ*ctpA* strain containing a plasmid encoding LmdC with a FLAG tag on the N- or C-terminus. Anti-LmdC immunoblot analysis of the input lysates revealed that the apparent size of the ∼ 12 kDa protein increased when the FLAG-tag was on the N-terminus, whereas the size of the ∼ 22 kDa protein increased when the tag was on the C-terminus (Fig. 1b). Anti-FLAG immunoprecipitation further confirmed that these proteins had the FLAG-tag (Fig. 1b). These data supported the hypothesis that a small N-terminal domain fragment of LmdC (LmdC^NTD^) accumulated in a Δ*prc* mutant, whereas a larger C-terminal domain fragment (LmdC^CTD^) accumulated in a Δ*ctpA* mutant.

### The N- and C-terminal domains of LmdC are separated by auto catalytic processing

We wanted to understand how the N- and C-terminal fragments of LmdC were generated. Cleavage by Prc and CtpA is not involved because the fragments were present in a Δ*prc* Δ*ctpA* strain (Fig. 1). To facilitate overproduction, the antigen we had used to generate our LmdC polyclonal antiserum was an LmdC-His_6_ protein lacking its predicted short N-terminal cytoplasmic domain and transmembrane helix (amino acids 1-46). When this protein was purified from the *E. coli* cytoplasm using nickel agarose, we isolated a band corresponding in size to the full-length protein (LmdC^FL^) as well as a smaller protein (Fig. 2a). The isolation of the smaller protein with nickel-agarose suggested that it retained the C-terminal His_6_ tag, and its size was similar to the C-terminal domain that accumulated in *P. aeruginosa*. This suggested that processing of LmdC to release its C-terminal domain had occurred, even though LmdC had been synthesized in the cytoplasm of *E. coli*. LmdC is a predicted LytM/M23 family endopeptidase and so we hypothesized that its own endopeptidase activity might be responsible for auto catalytic cleavage into its N- and C-terminal domains (NTD and CTD).

**FIG 2.**
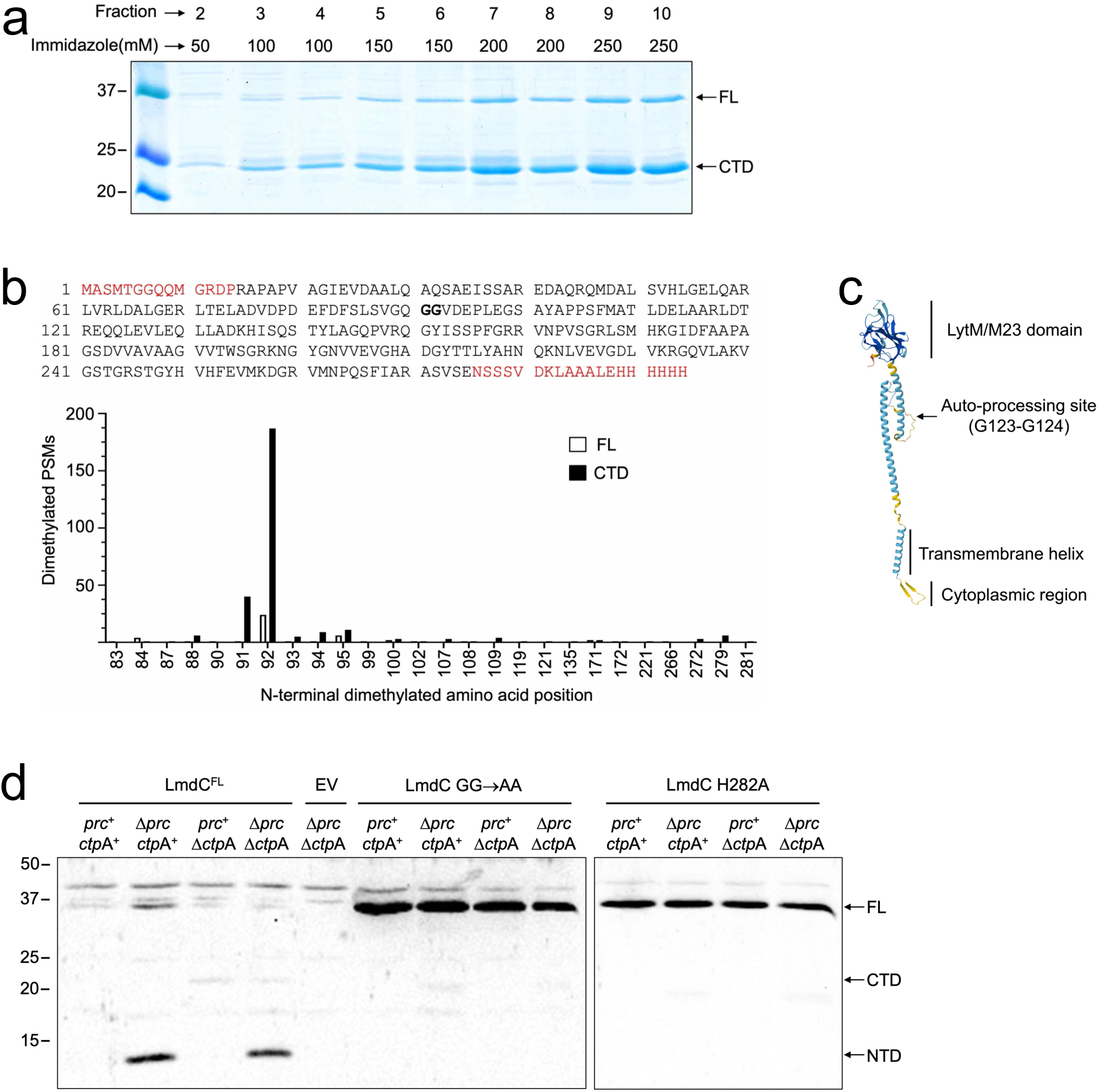
Evidence that the N- and C-terminal fragments of LmdC are generated by auto catalysis. (a) Elution of ′LmdC-His_6_ from NTA-agarose after production in *E. coli*. Fraction numbers and imidazole concentration in each one at the top, approximate sizes of molecular-mass marker proteins in kilodaltons on the left side. FL, full length; CTD; C-terminal domain. (b) Top: Sequence of the recombinant ′LmdC-His6 protein produced in *E. coli*, which includes vector-derived amino acids in red and LmdC without its N-terminal cytoplasmic and transmembrane domains in black. Bold GG at positions 91 and 92 are the predicted auto processing site, which corresponds to positions 123 and 124 in wild type LmdC. Bottom: Graph summarizing mass spectrometry identification of N-terminal demethylated peptides (see text) from LmdC FL and CTD. Amino acid positions correspond to the protein sequence at the top. PSMs, peptide spectral matches. (c) AlphaFold-predicted structure of LmdC with the N-terminus at the bottom. Model confidence, dark blue = very high, cyan = high, yellow = low, orange = very low. (d) Anti-LmdC immunoblot of whole-cell lysates of Δ*lmdC* strains with the indicated version of LmdC encoded on a plasmid or the empty vector (EV). GG→AA is the Gly-123 Gly-124 to Ala-123 Ala-124 mutation. Arrows on the right indicate LmdC full-length (FL), C-terminal domain (CTD) and N-terminal domain (NTD) protein bands. Approximate sizes of molecular-mass marker proteins in kilodaltons on the left. Single representative immunoblot of replicate experiments.

To determine the likely processing site, we took advantage of the purified LmdC^CTD^-His_6_ protein by characterizing its N-terminus. The LmdC^FL^-His_6_ and LmdC^CTD^-His_6_ proteins (Fig. 2a) were incubated with formaldehyde, which dimethylates N-terminal amino groups (as well as amino groups of lysine side chains). Trypsinized peptides were analyzed by mass spectrometry to detect those with a dimethylation mass increase of 28 Da. Gly-92 of the recombinant protein, which corresponds to Gly-124 of wild type LmdC, was the primary site of non-lysine dimethylation in the LmdC^CTD^-His_6_ protein, which identified this residue as its N-terminal amino acid (Fig. 2b; Supplemental Files 1-2). A small amount of dimethylation of this glycine was also detected for the LmdC^FL^-His_6_ protein, which our Proteomics core suggested was most likely due to carryover contamination because the LmdC^CTD^-His_6_ protein was analyzed first. However, this does not alter the main conclusion that these data suggested that LmdC is cleaved between amino acids Gly-123 and Gly-124 to generate the NTD and CTD fragments (Fig. 2b). In the AlphaFold predicted structure of LmdC, these glycines are in an unstructured region between two periplasmic alpha helices, which could make the bond between them easily accessible for proteolysis (Fig. 2c).

To test if Gly-123 and Gly-124 were required for LmdC processing in *P. aeruginosa,* and if the processing was likely to be auto catalytic, we analyzed two mutant LmdC proteins. The LytM/M23 family of metallopeptidases is characterized by zinc-dependence and a HxH motif (32). The first histidine in this motif is predicted to coordinate and activate an incoming water molecule, which triggers the nucleophilic attack on the scissile bond, whereas the second histidine is involved in zinc ion coordination (33, 34). A previous study suggested that mutating the zinc-coordinating histidine in the HxH motif might misfold the protein, whereas mutating the catalytic histidine might not (35). Therefore, to destroy LmdC activity we mutated the predicted catalytic histidine to alanine (H282A). Separately, we also mutated Gly-123 and Gly-124 flanking the predicted processing site to alanines (GG→AA). Plasmids encoding LmdC, LmdC^H282A^ and LmdC^GG→AA^ were put into Δ*lmdC* strains with various *prc*/*ctpA* genotypes and analyzed by anti-LmdC immunoblot. As expected, with LmdC we observed the accumulation of the ∼12 kDa NTD in a Δ*prc* mutant, the ∼22 kDa CTD in a Δ*ctpA* mutant, and both domains in a Δ*prc* Δ*ctpA* mutant (Fig. 2d; note that the polyclonal serum recognizes the CTD poorly and so it was only faintly detected). However, for both LmdC^H282A^ and LmdC^GG→AA^, regardless of the *prc* or *ctpA* genotype, we only detected a larger protein corresponding in size to unprocessed LmdC (Fig. 2d). These data support the identification of Gly-123 -Gly-124 as the processing site and the hypothesis that the processing is auto catalytic. Together with the data in Figure 1, they also show that unprocessed LmdC^FL^ did not accumulate in Δ*prc* or Δ*ctpA* mutants, suggesting that LmdC is not degraded by either protease prior to being auto processed.

### *In vitro* proteolysis of LmdC^NTD^ by Prc

We already showed previously that what we now know to be the CTD of LmdC is degraded by CtpA *in vitro*, consistent with its accumulation in Δ*ctpA* strains (14). To test if the NTD of LmdC is a direct substrate of Prc, we constructed a plasmid encoding a maltose binding protein (MBP)-LmdC^NTD^ fusion protein and purified it with amylose resin, which was the approach we used to confirm MucA22 periplasmic domain degradation by Prc (30). Prc-His_6_ and Prc-S456A-His_6_ (catalytic mutant) proteins were purified from the periplasm of an *E. coli ΔnlpI* and *Δprc::kan* mutant to avoid contamination with *E. coli* Prc or its accessory protein NlpI (13). When incubated together, we found that Prc degraded MBP-LmdC^NTD^ whereas the Prc-S456A negative control did not (Fig. 3a). The MBP-LmdC^NTD^ fusion protein was not fully degraded, with the size of the reaction product consistent with degradation of LmdC^NTD^ but not MBP. This was supported by anti-LmdC and anti-MBP immunoblot analysis, and the same phenomenon had occurred previously with a MBP-′MucA22 protein (Fig. 3a) (30). As an additional specificity control, we tested if Prc degraded a His_6_-LmdC^CTD^ protein, but it did not, which is consistent with LmdC^CTD^ failing to accumulate in a Δ*prc* mutant (Fig. 3b). These data support the conclusion that Prc degrades the NTD of LmdC specifically, which is consistent with the accumulation of only this part of LmdC in Δ*prc* strains.

**FIG 3.**
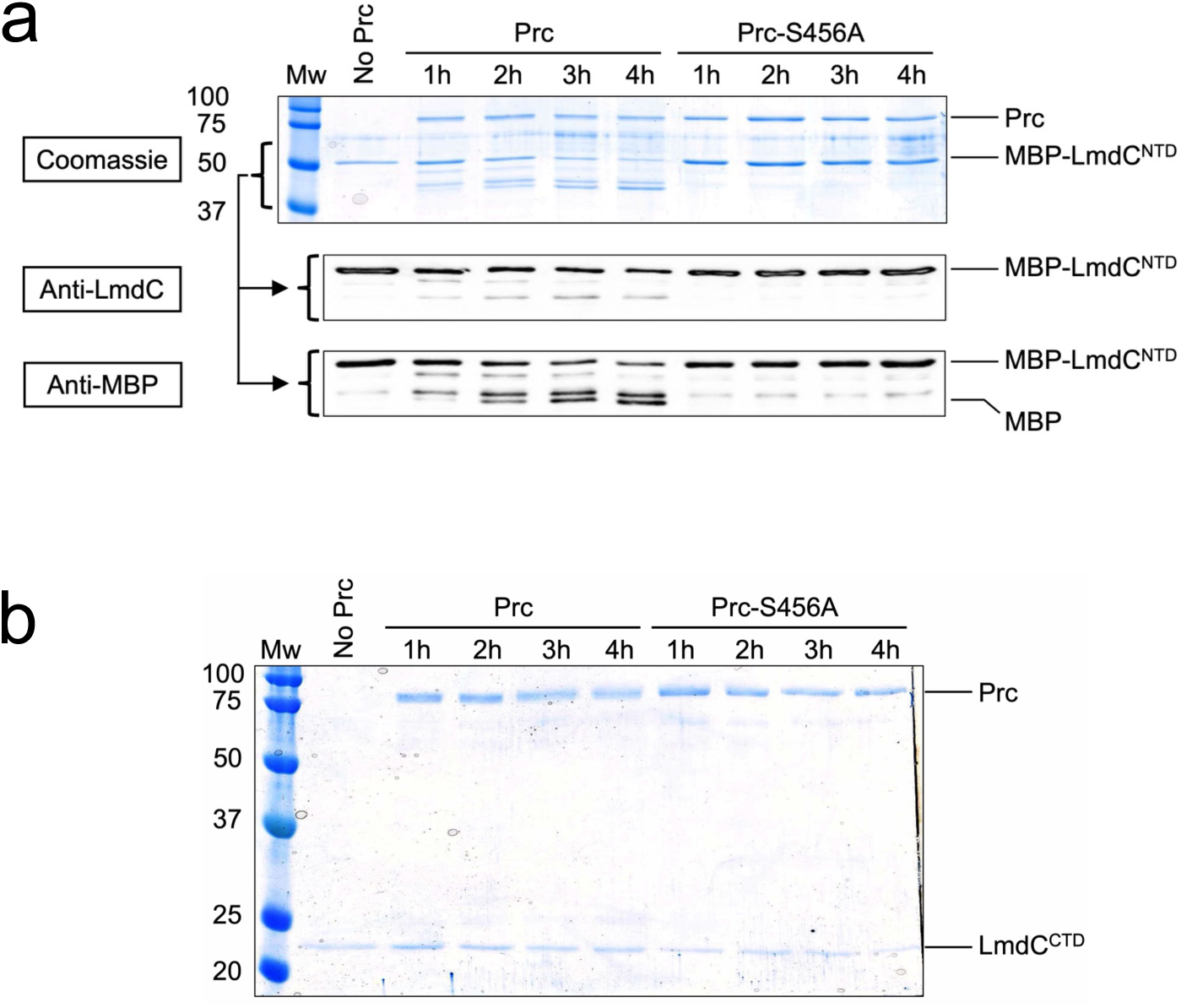
Analysis of LmdC domain cleavage by Prc *in vitro*. (a) MBP-′LmdC^NTD^ protein was incubated with Prc-His6 or Prc-S456A-His6 proteins at 37°C for the indicated number of hours. The top image shows a Coomassie blue-stained SDS-PAGE gel with molecular-mass marker proteins in kilodaltons to the left (Mw). The middle and bottom images are anti-LmdC and -MBP immunoblots, with the bracket showing the corresponding region of the SDS-PAGE gel. (b) LmdC^CTD^-His6 protein was incubated with Prc-His6 or Prc-S456-His6 proteins at 37°C for the indicated number of hours and analyzed by SDS-PAGE followed by Coomassie blue-staining. Molecular-mass marker proteins in kilodaltons to the left (Mw). Data presented are single representatives of replicate experiments.

### Analysis of LmdC localization *in vivo*

CtpA associates with outer membrane lipoprotein LbcA and so at least part of the reason it might be unable to degrade LmdC^FL^ and LmdC^NTD^ is their predicted inner membrane location. In contrast, the LmdC^CTD^ could be released as a soluble protein in the periplasm, facilitating its degradation by CtpA. In *E. coli*, Prc also associates with an outer membrane lipoprotein, NlpI (13). However, there is no NlpI homolog in *P. aeruginosa*, and we have no evidence that Prc works with an accessory outer membrane lipoprotein. This could leave Prc free to access the exposed periplasmic region of LmdC^NTD^ anchored in the inner membrane. To investigate this, we analyzed protein localizations by immunoblot in strains with various *prc*, *ctpA* and *lmdC* genotypes. Prc was located entirely in the soluble fraction, consistent with it being unassociated with a lipoprotein partner (Fig. 4a). Analysis of cells with the *lmdC* GG→AA auto-processing mutation showed that LmdC^FL^ was also in the membrane fraction (Fig. 4b). As predicted, in *lmdC*^+^ cells where most LmdC is auto processed into its NTD and CTD, the CTD was in the soluble fraction and the NTD was in the membrane fraction (Fig. 4b). Finally, we fractionated membranes on discontinuous sucrose gradients to more precisely locate LmdC^FL^ and LmdC^NTD^, and both fractionated similarly to inner membrane control protein LepB (signal peptidase I), supporting an integral inner membrane location (Fig. 4c).

**FIG 4.**
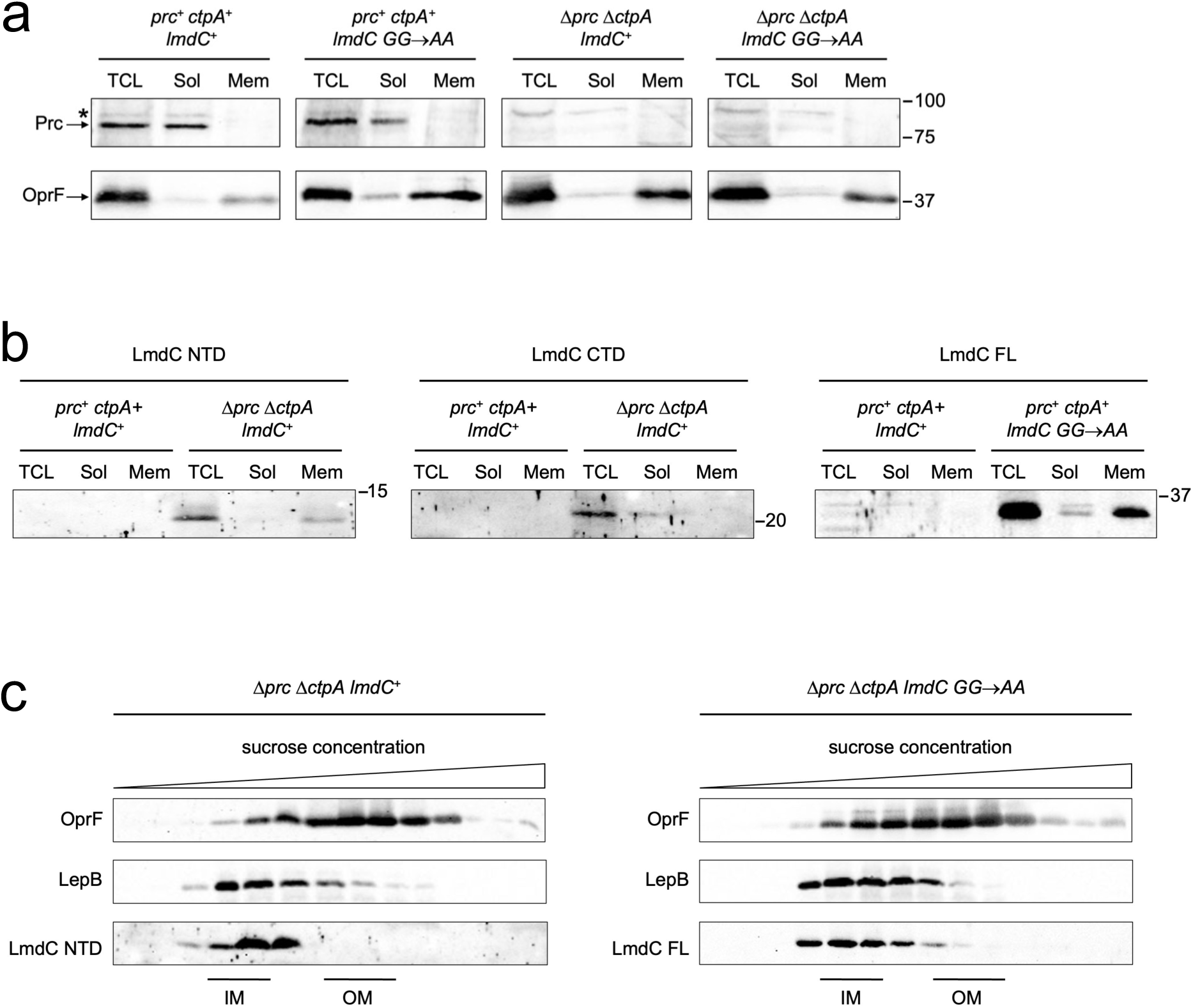
Prc and LmdC localization analysis. (a) Anti-Prc and anti-OprF (membrane control) immunoblot analysis of membrane and soluble fractions from strains with the genotypes at the top. TCL = total cell lysate; Sol = soluble fraction; Mem = membrane fraction. Approximate sizes of molecular-mass marker proteins (M), in kilodaltons, are on the right. The asterisk indicates a protein cross-reactive with the Prc polyclonal antiserum. (b) Anti LmdC immunoblot analysis using subcellular fractions samples in panel a. The form of LmdC detected is shown at the top, N-terminal domain (NTD), C-terminal domain (CTD) and full-length (FL). Approximate sizes of molecular-mass marker proteins (M), in kilodaltons, are on the right. The *prc*^+^ *ctpA*^+^ *lmdC*^+^ samples served as negative controls because the levels of LmdC FL, NTD and CTD were too low for detection due to auto processing and degradation. (c) Membrane fractionation analysis. Membranes from strains with the indicated genotypes were fractionated on discontinuous sucrose gradients as described in Materials and Methods. Fractions 7-20 were analyzed by immunoblotting with antisera to detect LmdC, OprF (outer membrane control) or LepB (inner membrane control). Approximate locations of peak fractions for the inner (IM) and outer (OM) membranes based on the control proteins are indicated. For all panels GG→AA is the Gly-123 Gly-124 to Ala-123 Ala-124 auto-processing defect mutation of *lmdC*.

### Evidence that LmdC auto processing is detrimental to its function *in vivo*

The activity of *Staphylococcus aureus* LytM increases *in vitro* following removal of its N-terminal domain, although the *in vivo* significance of this is unclear (35). Some other LytM/M23 family members have inhibitory N-terminal pro-peptides (33). Therefore, LmdC^FL^ could be a precursor protein, with the NTD inhibiting the function of the catalytic CTD. An alternative possibility is that LmdC^FL^ is the functional enzyme, with auto processing serving only to initiate its proteolytic destruction by Prc and CtpA. Experiments to distinguish between those possibilities required an LmdC-associated phenotype, but individual autolysin mutants often lack robust phenotypes due to redundancy in the laboratory. We did find that a Δ*lmdC* mutation decreased resistance to ticarcillin/clavulanate (TIC), a β-lactam/β-lactamase inhibitor combination used in the clinic, but the phenotype was too modest for robust genetic analyses: E-test strip minimal inhibitory combination (MIC) 8 µg/ml (*lmdC*^+^) vs. 6 µg/ml (Δ*lmdC*). However, introducing a Δ*lmdC* mutation into a Δ*mepM* strain caused an 8-fold decrease in TIC resistance: E-test MIC 16 µg/ml (Δ*mepM*) vs. 2 µg/ml (Δ*mepM* Δ*lmdC*). MepM is another LytM/M23 enzyme degraded by CtpA (14). A growth curve turbidity assay in the presence of TIC provided us with a robust and sensitive Δ*lmdC* phenotype in a Δ*mepM* strain, characterized by near normal increase in turbidity in exponential phase followed by a rapid decrease in stationary phase (Fig 5a).

**FIG 5.**
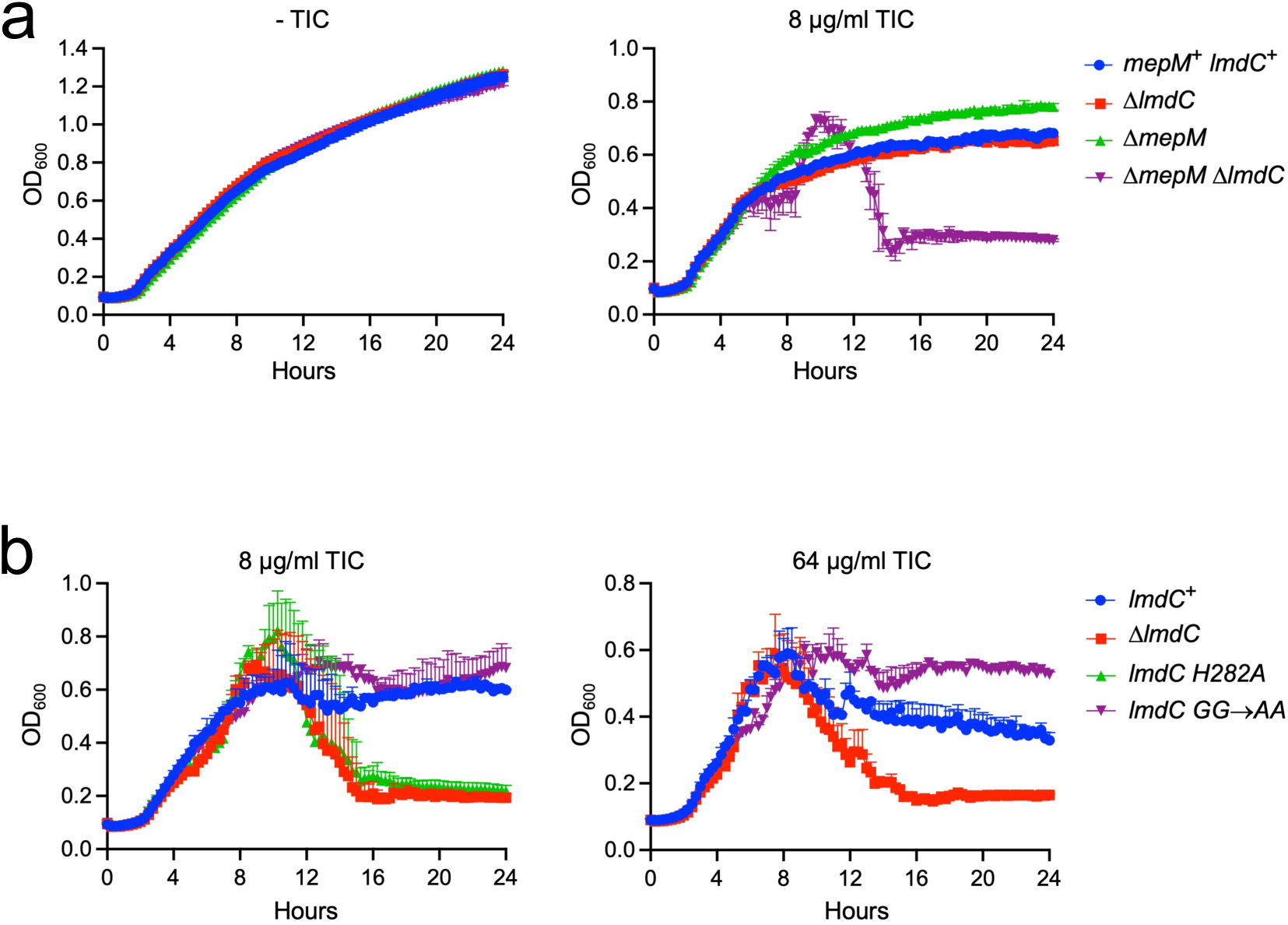
Growth phenotypes in the presence of ticarcillin-clavulanate suggest auto processing is detrimental to LmdC function. (a) Effects of Δ*mepM* and Δ*lmdC* mutations on resistance to ticarcillin-clavulanate. Strains were grown in 96-well plates at 37°C in LB broth containing 8 µg/mL of ticarcillin-clavulanate 15:1 (TIC) or LB broth only (-TIC). Optical density at 600nm was measured every 30 minutes in a Bioscreen 6 device for 24 hours. Data are the average of duplicate cultures with error bars showing the range. (b) Δ*mepM* strains with the indicated *lmdC* genotypes were grown as in panel a with 8 µg/mL or 64 µg/mL TIC. Data are the average of triplicate cultures with error bars showing only the positive standard deviation to improve clarity. GG→AA is the Gly-123 Gly-124 to Ala-123 Ala-124 auto processing defect mutation of *lmdC*. The H282A mutation affects a predicted catalytic histidine of LmdC.

We constructed Δ*mepM* strains with one of three different mutations of the chromosomal *lmdC* gene: a complete deletion (Δ*lmdC*), the catalytically inactive mutant (LmdC^H282A^) and the auto processing defective mutant (LmdC^GG→AA^). The H282A mutation phenocopied the Δ*lmdC* mutation, as expected, but the auto processing mutant had a *lmdC*^+^ phenotype with 8 µg/ml TIC (Fig. 5b). Furthermore, when we increased the TIC concentration to 64 µg/ml, the auto processing mutant maintained a higher turbidity in stationary phase than the *lmdC*^+^ strain (Fig. 5b). These data suggest that auto processing is detrimental to LmdC function. This supports the hypothesis that LmdC^FL^ is the functional endopeptidase with auto processing serving only to trigger its destruction by Prc and CtpA.

### A phenotypic link between LmdC, bactofilin and PA0663

Many *lmdC* homologs are linked to a gene encoding bactofilin, a polymer-forming cytoskeletal protein that recruits PG enzymes for morphogenesis, cell division, or motility (e.g., 36, 37-39). In two Alphaproteobacteria, LmdC and bactofilin proteins co-localized, and the conserved β-hairpin-containing cytoplasmic domain of *Hyphomonas neptunium* LmdC bound to bactofilin *in vitro* (Fig. 2c) (39). *P. aeruginosa* has one gene encoding an uncharacterized bactofilin domain protein (PA0664, hereafter called *baf*), but it is not present in most *Pseudomonas* species. However, all *Pseudomonas* species with *baf* also have *lmdC*, and all species with *lmdC* also have *baf* (40). This absolute co-occurrence of *lmdC* and *baf* in *Pseudomonas* species suggested the likelihood of a functional link between them. In the *lmdC*^+^ species, *lmdC* occurs in two different locations: either in the same predicted operon as *baf* or adjacent to a gene essential for DNA replication (*dci*) and the division and cell wall (*dcw*) cluster (Fig. 6a). For those species where *lmdC* and *baf* are not linked, which includes *P. aeruginosa*, *baf* is in a predicted operon with gene of unknown function PA0663. All these observations prompted us to investigate if there is a link between LmdC, bactofilin and PA0663. We will refer to PA0663 as Api (<u>a</u>uto <u>p</u>rocessing inhibitor) due to subsequent experiments implicating its involvement together with Baf in inhibiting LmdC auto processing.

**FIG 6.**
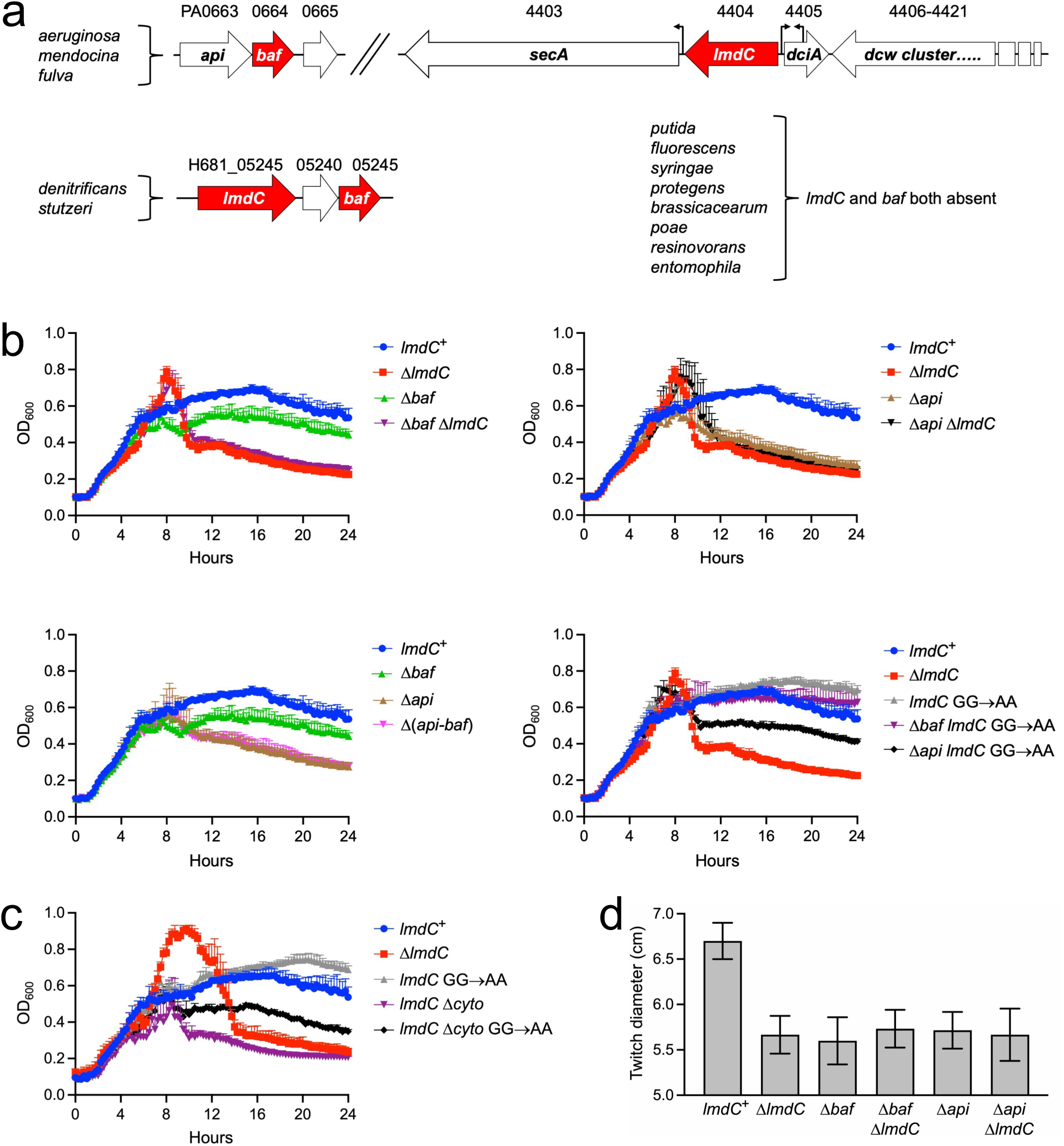
Evidence of a functional link between LmdC, Baf and Api. (a) Occurrence and arrangement of *lmdC* and *baf* genes in *Pseudomonas* species (from the *Pseudomonas* ortholog database). Genes from *P. aeruginosa* strain PAO1 and *P. denitrificans* ATCC 13867 are shown as examples of the two arrangements. *P. denitrificans* ATCC 13867 is also now known as *P.* sp. ATCC 13867 and recent phylogenetic studies reclassified *stutzeri* into a separate *Stutzerimonas* genus. (b) Δ*mepM* strains with the indicated *lmdC, baf and api* genotypes were grown in 96-well plates at 37°C in LB broth containing 64 µg/mL of ticarcillin-clavulanate 15:1. Optical density at 600nm was measured every 30 minutes in a Bioscreen 6 device for 24 hours. All strains/cultures in panel b were grown simultaneously but are shown in separate graphs for clarity, with some data duplicated to facilitate comparisons (for example, all four graphs shown the same *lmdC*^+^ strain data). (c) Δ*mepM* strains with the indicated *lmdC* genotypes were grown as in panel b. For panels b and c data are the average of triplicate cultures with error bars showing only the positive standard deviation to improve clarity. GG→AA is the Gly-123 Gly-124 to Ala-123 Ala-124 auto processing defect mutation of *lmdC*. The Δ*cyto* mutation removes the predicted cytoplasmic domain of LmdC. In media without ticarcillin-clavulanate the growth curves of all strains in panels b and c were indistinguishable (Fig. 5 and not shown). (d) Diameter of zone of twitching motility of strains with the indicated genotypes, determined as described in Materials and Methods. Data are the average of triplicate assays with error bars showing the standard deviations from the mean. For all five deletion mutants *P* < 0.01 in comparison to the *lmdC*^+^ strain (unpaired t test).

We constructed Δ*mepM* strains with combinations of *lmdC*, *baf* and *api* mutations and used the LmdC-associated TIC-resistance phenotype to investigate the impact of Baf and Api. Like a Δ*lmdC* mutation, a Δ*baf* mutation decreased turbidity in stationary phase, but the magnitude was less severe (Fig. 6b). This suggested that Baf might assist LmdC function but is not essential for LmdC function. Two additional observations were consistent with that suggestion: (1) Δ*lmdC* was epistatic to Δ*baf* and (2) Δ*baf* reduced the higher stationary phase turbidity phenotype of the *lmdC*^GG→AA^ auto processing defective mutant (Fig. 6b). A *Δapi* mutant phenocopied a Δ*lmdC* mutant. Additionally, the Δ*api* and Δ*lmdC* mutations were epistatic to each other, and Δ*api* was also epistatic to Δ*baf* (Fig. 6b). These observations are consistent with Api playing one or more essential roles in LmdC function. We considered two possibilities, which are not mutually exclusive: (1) Api might assist the endopeptidase function of LmdC^FL^ and/or (2) Api might inhibit LmdC auto processing. The *lmdC*^GG→AA^ mutation, which eliminates auto processing, was not epistatic to Δ*api*. This supports the possibility that Api assists LmdC^FL^ function independently of influencing auto processing (Fig. 6b). However, it did not rule out the possibility that Api might also inhibit auto processing of the wild LmdC protein, and our later experiments suggested that it does.

The β-hairpin-containing cytoplasmic domain of *H. neptunium* LmdC bound to bactofilin *in vitro* (39). This cytoplasmic β-hairpin is predicted to be widely conserved in the LmdC clade, including in *P. aeruginosa* LmdC (31) (Fig. 2c). Therefore, we tested the effect of an in-frame deletion mutation that removed this region (*lmdC*^Δcyto^) and found that it decreased stationary phase turbidity similarly to a Δ*lmdC* mutation (Fig. 6c). It also reduced stationary phase turbidity in the *lmdC*^GG→AA^ auto processing defective mutant. These observations are consistent with an important functional role for the conserved cytoplasmic domain of LmdC in *P. aeruginosa*, which may involve communication with Baf and/or the cytoplasmic domain of Api (see Discussion).

### Twitching motility phenotypes are also consistent with functional links between LmdC, bactofilin and Api

Expression of *lmdC* and *baf* is positively regulated by cAMP and the Vfr regulator, which control cell surface functions, including the type IV pilus (T4P) (41, 42). For the T4P, Pseudomonads have been divided into “simple” and “complex” species, with the latter having a more complex *chp* gene cluster that controls T4P-dependent twitching motility (43, 44). There are 59 genes present in all complex species that are absent from all simple species, and several of these genes are in the Vfr regulon (43). They include *lmdC* and *baf*, the additional genes in the *chp* cluster, the adenylate cyclase gene *cyaB*, and T4P regulators *fimL* and *fimX* (43, 45, 46). These findings raised the possibility that LmdC might influence the T4P and so we tested the effects of *lmdC*, *baf* and *api* null mutations on twitching motility. All three mutations reduced the diameter of the zone of twitching motility (Fig. 6d). Although the effects were modest, they were comparable to those of some other mutations that influence but do not abolish T4P function (47). Also, as with the TIC-sensitivity phenotype, Δ*lmdC* was epistatic to Δ*baf* and Δ*api*. We cannot conclude from these data that LmdC plays a direct role in T4P function. Even so, regardless of the underlying mechanism it provided a second independent assay in which Δ*lmdC*, Δ*baf* and Δ*api* mutations caused similar phenotypes, further supporting functional links between them.

### *api*-*baf* overexpression stabilizes LmdC^FL^

LmdC auto processing is likely to be controlled rather than constitutive if it serves only to trigger LmdC destruction by Prc and CtpA. Our experiments had implicated Baf and Api in facilitating the function of LmdC^FL^, including the LmdC^GG→AA^ version that cannot auto process. One possibility is that Baf and Api promote a location-specific LmdC^FL^ role. In this scenario, any LmdC^FL^ that escapes the influence of Baf and Api could cleave peptidoglycan in the wrong place, but its immediate auto processing and degradation would defend against this. This scenario predicts that the presence of Baf and/or Api should reduce LmdC auto processing. Our previous immunoblot experiments had suggested that the majority of LmdC in the cell is already auto processed, making attempts to detect any further increased auto processing in Δ*baf* and Δ*api* mutants difficult (Figs. 1a and 2d). Therefore, as an alternative approach to test if Baf and/or Api might inhibit LmdC auto processing, we tested if *baf* and/or *api* overexpression stabilized LmdC^FL^.

In a *lmdC^+^* strain, plasmids encoding Api or Baf alone had no obvious effect on the low level of detectable LmdC^FL^, but when *api*-*baf* were both overexpressed, there was an unequivocal increase in the amount of LmdC^FL^ (Fig. 7a). This is consistent with *api*-*baf* overexpression reducing LmdC auto processing. In further support of that conclusion, *api*-*baf* overexpression did not affect the amount of LmdC^FL^ with the GG→AA mutation, which cannot auto process (Fig. 7a). *api*-*baf* overexpression also had no effect on the level of LmdC^FL^ without its cytoplasmic domain (*lmdC*^Δcyto^), which provides further support for the importance of this conserved domain (Fig. 7a). The phenotypes in the *lmdC*^GG→AA^ and *lmdC*^Δcyto^ strains also demonstrated that the elevated level of LmdC^FL^ caused by *api*-*baf* overexpression was not to be due to increased *lmdC* gene expression because all three alleles of *lmdC* were at the native chromosomal locus.

**FIG 7.**
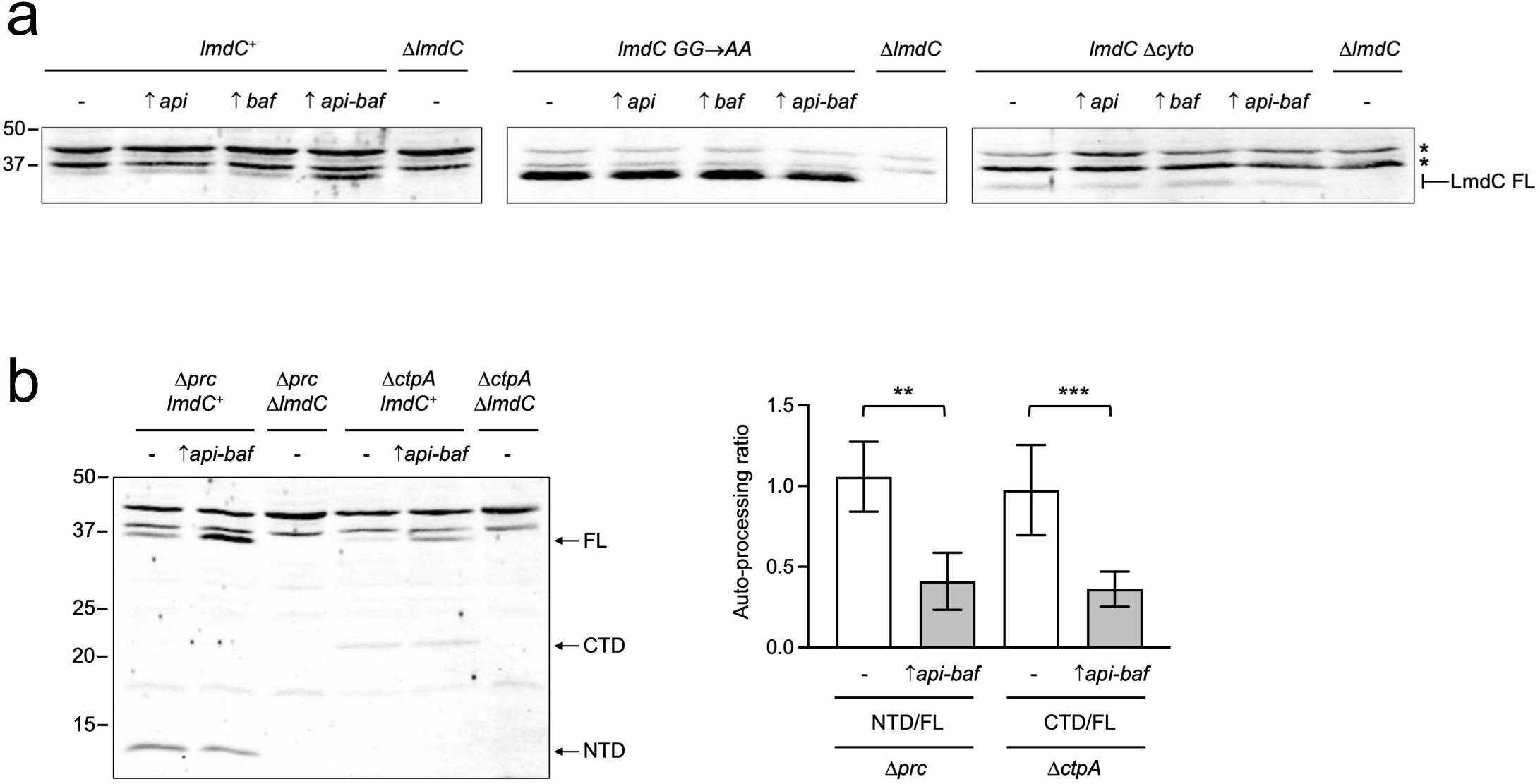
Overexpression of *api-baf* inhibits LmdC auto processing. (a) Effect of *api* and/or *baf* overexpression on the level of full length LmdC. Anti-LmdC immunoblot of whole-cell lysates of strains with the indicated allele of *lmdC* at the native chromosomal locus. GG→AA is the Gly-123 Gly-124 to Ala-123 Ala-124 auto processing defect mutation. The Δ*cyto* mutation removes the predicted cytoplasmic domain. Strains contained an *araBp* expression plasmid to overexpress the indicated gene(s) or the empty vector (-). Asterisks indicate two higher molecular weight proteins cross-reactive with the polyclonal LmdC antiserum. The location of the full length (FL) LmdC protein is indicated on the right, with LmdC Δcyto version running slightly lower than the other versions due to the truncation. Approximate sizes of molecular-mass marker proteins in kilodaltons on the left. Data is a single representative of replicate experiments. (b) Effect of *api-baf* overexpression on LmdC auto processing. Anti-LmdC immunoblot of whole-cell lysates of strains with the indicated genotypes containing an *araBp* expression plasmid overexpressing *api-baf* or the empty vector (-). Arrows on the right indicate LmdC full-length (FL), C-terminal domain (CTD) and N-terminal domain (NTD) protein bands. Approximate sizes of molecular-mass marker proteins in kilodaltons on the left. A Single representative immunoblot is shown on the left. The graph on the right is the mean data from 4-6 independent immunoblots, including the one on the left, showing the LmdC NTD/full length (FL) or CTD/FL ratios +/-SD determined by quantitative immunoblot analysis. **P < 0.01; ***P < 0.001 (unpaired t test).

Degradation of the LmdC^NTD^ and LmdC^CTD^ fragments by CtpA and Prc in the preceding experiment meant that their levels could not be monitored to further assess effects on auto processing. We had not used a Δ*prc* Δ*ctpA* strain background because we were concerned that removal of these proteases would alter cell envelope physiology significantly, which might impact processes involved in controlling LmdC auto processing. A Δ*ctpA* mutant is known to have significant effects on cell envelope-associated processes, including type III secretion and surface attachment (14, 48). Indeed, for unknown reasons we found that LmdC^FL^ was less abundant in a Δ*ctpA* mutant and difficult to detect in a Δ*prc* Δ*ctpA* double mutant (Fig. 7b and not shown). Nevertheless, with this caveat in mind, and attempting to minimize any impact, we did experiments in single protease defective strains. This was feasible because only one of the processed LmdC^NTD^ and LmdC^CTD^ fragments needs to be detected to follow auto processing. Therefore, we monitored the LmdC^NTD^/LmdC^FL^ ratio in a Δ*prc* strain, where LmdC^NTD^ is not degraded, and the LmdC^CTD^/LmdC^FL^ ratio in a Δ*ctpA* strain, where LmdC^CTD^ is not degraded. Quantification showed that *api*-*baf* overexpression caused a similar reduction in the LmdC^NTD^/LmdC^FL^ and LmdC^CTD^/LmdC^FL^ ratios (Fig. 7b). This supports the conclusion that the effect of Api-Baf overproduction on the level of LmdC^FL^ is due to reduced LmdC auto processing. The observation that *api-baf* overexpression affected auto processing to the same extent in Δ*prc* and Δ*ctpA* mutants provides increased confidence because the same conclusion was reached from experiments in these two independent strains.

## DISCUSSION

Based on data presented here, we propose a working model for regulated auto catalytic processing promoting autolysin destruction by two proteases (Fig. 8a). There are some limitations in our study, which include reliance on overexpression in the final experiment and a lack of protein-protein interaction data, as discussed later. Therefore, our model also includes hypotheses to help guide future experiments. Baf and Api might promote a restricted peptidoglycan cleavage role for LmdC, perhaps at a specific location. Baf could act as an anchor to bring Api and LmdC together at the site of action, consistent with a bactofilin location-specific anchor role in other species (36–39). For Api, our data support the possibility of two complimentary roles to promote LmdC function. First, inhibiting LmdC auto processing, supported by the data in Figure 7. Perhaps this could be achieved by a coiled-coil interaction between the periplasmic alpha helices of Api and LmdC preventing the LmdC C-terminal catalytic domain accessing its Gly-Gly auto processing site (Fig. 8). Second, the detrimental effect of a Δ*api* mutation in a *lmdC*-GG→AA mutant, in which LmdC cannot auto process, suggests that Api also facilitates the function of full length LmdC (Fig. 6b). This might be achieved by the C-terminal domain of Api anchoring to the cell wall target of LmdC or regulating LmdC activity by some other means (Fig. 8a). Implicit in our model is that any LmdC that goes rogue by escaping Api and Baf could cleave the cell wall at the wrong location, but this is defended against by its immediate auto processing and degradation by Prc and CtpA (Fig. 8a). Our model also proposes explanations for the inability of Prc and LbcA•CtpA to target unprocessed LmdC for degradation. Its integral inner membrane location might make it difficult for the outer membrane-anchored LbcA•CtpA complex to access, and the C-terminus of full length LmdC is probably not recognized by Prc, in contrast to the predicted unstructured C-terminus of LmdC^NTD^ (Fig. 3).

**FIG 8.**
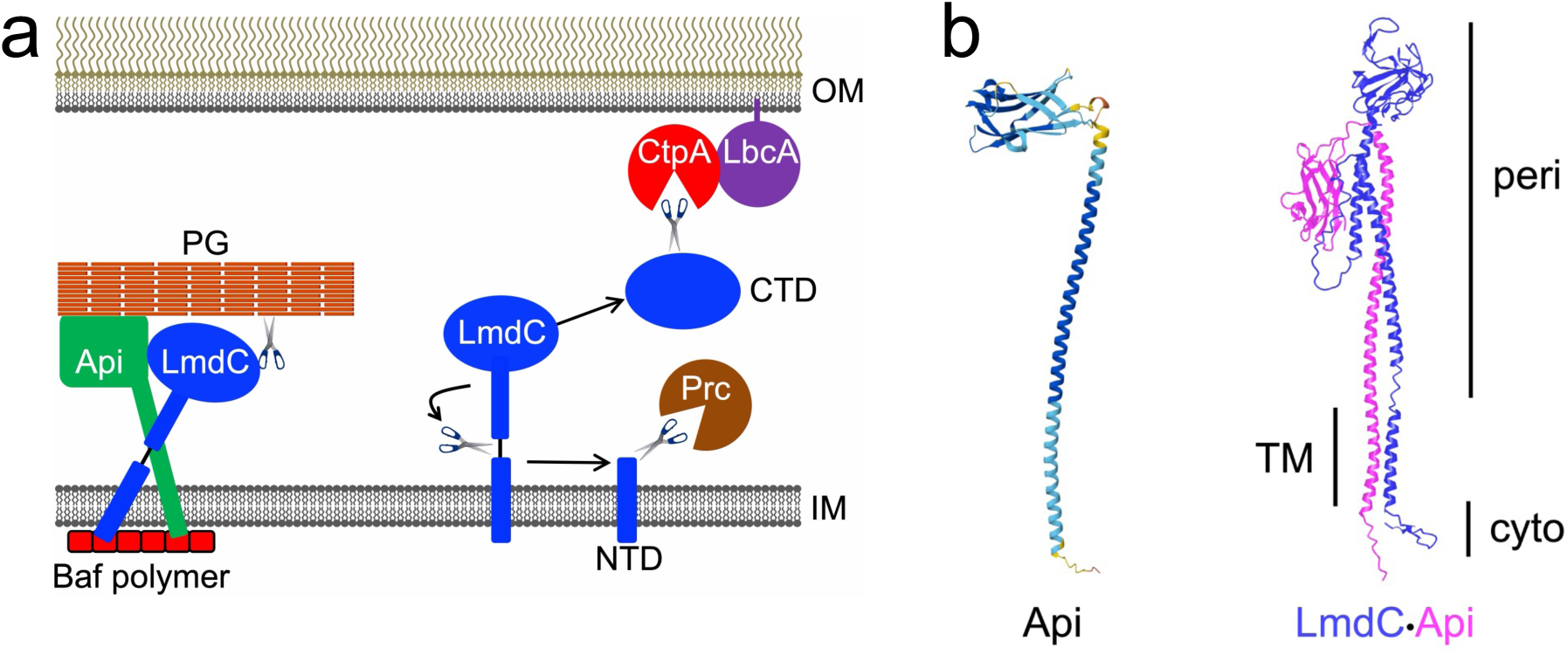
Working model of the LmdC control mechanism. (a) A hypothetical trimolecular Baf-Api-LmdC association promotes a role of LmdC in peptidoglycan (PG) cleavage. Baf might ensure this occurs at a specific location and bring Api and LmdC together by anchoring their short cytoplasmic domains. A coiled-coil interaction between the periplasmic alpha helices of LmdC and Api could ensure that the catalytic domain of LmdC can only access PG and not its auto processing site, in *cis* or in *trans*. The C-terminal globular domain of Api might anchor the complex to its PG target or interact with the catalytic domain of LmdC to control its activity. To prevent dangerous PG cleavage at the wrong location, any free LmdC is immediately auto processed into N- and C-terminal domains (NTD, CTD) that are degraded by Prc and the LbcA•CtpA complex. OM = outer membrane, IM = inner membrane. (b) AlphaFold-predicted structure of Api on the left with the N-terminus at the bottom. Model confidence, dark blue = very high, cyan = high, yellow = low, orange = very low. On the right is the highest ranked AlphaFold Multimer model of a LmdC•Api complex. peri = periplasmic region, TM = transmembrane region, cyto = cytoplasmic region.

LmdC is a LytM/M23 family endopeptidase. The original LytM of *Staphylococcus aureus* is a glycylglycine endopeptidase and several other LytM/M23 enzymes cleave bonds that involve at least one glycine (32, 49). This fits with LmdC auto processing occurring by cleavage of a Gly-Gly bond (Fig. 2). Some individual LytM/M23 family members have broader peptide bond specificity that includes bonds with or without glycine, which could allow LmdC to cleave both its own Gly-Gly bond and the stem peptide cross-links of *P. aeruginosa* peptidoglycan, which do not involve glycine (14, 33). The LmdC clade of LytM/M23 endopeptidases is also widely conserved (31). These proteins typically feature a short cytoplasmic tail, a transmembrane domain, periplasmic alpha helices and a C-terminal peptidase domain. AlphaFold predictions suggest that most of the 31 LmdC proteins that were analyzed to define the LmdC clade also have a Gly-Gly bond in a predicted unstructured region between alpha helices (31, 50). Others have Ala-Gly or Met-Gly bonds in this region, which might also be auto processed if both glycines are not required. Therefore, LmdC auto processing is unlikely to be limited to *Pseudomonas aeruginosa*. In fact, previous studies reported difficulty constructing a stable *H. neptunium* LmdC fluorescent fusion protein and observed only low-level diffuse fluorescence from an *A. tumefaciens* LmdC fluorescent fusion protein. In both cases LmdC was tagged at the C-terminus, suggesting that these findings could be explained by LmdC auto-processing.

Bactofilins are polymer-forming cytoskeletal proteins that interact with the inner face of the cytoplasmic membrane, usually at regions of high curvature (51–55). They promote site-specific localization of proteins involved in cell wall biology, including some autolysins (39, 51, 53–56). Therefore, our working model has Baf acting as a location specific anchor for LmdC (Fig. 8a). Bactofilin genes are often closely linked to genes encoding LytM/M23 endopeptidases, including members of the LmdC clade, suggesting a common functional link between members of these protein families (39). In *Helicobacter pylori*, a bactofilin interacts with an LmdC-containing complex to promote localized peptidoglycan synthesis and the characteristic cell shape (55, 57). In the Alphaproteobacterial species *H. neptunium* and *Rhodospirillum rubrum*, a bactofilin is required for the site-specific localization of LmdC proteins to facilitate precise editing of peptidoglycan for stalk formation in *H. neptunium* and cell curvature in *R. rubrum* (39). If Baf does aid localization of LmdC in *P. aeruginosa,* the intermediate TIC-sensitivity phenotype of a Δ*baf* mutant might indicate that at least some LmdC can still function without Baf, but with reduced efficiency (Fig. 6b). *P. aeruginosa* has a simple rod-shaped cell, does not undergo asymmetric cell division, and is not known to exhibit complex morphological changes of individual cells. However, possibilities for a LmdC location-specific role could be related to cell division, T4P function, or both. The latter is plausible because *P. aeruginosa* T4P components are recruited to future division sites to facilitate preinstallation of the T4P into nascent poles (58). Consistent with these possibilities are the observations that in some *Pseudomonas* species *lmdC* and the division and cell wall (*dcw*) cluster are linked (Fig. 6a); that *lmdC* is only present in *Pseudomonas* species with a more complex *chp* gene cluster that controls T4P-dependent twitching motility (43, 44); and that Δ*lmdC*, Δ*baf* and Δ*api* mutations all reduced T4P-dependent twitching motility (Fig. 6d). It is also notable that a *Myxococcus xanthus* bactofilin is involved in polar localization of two T4P motor ATPases (59).

The Genomic Context Visualizer (GeCoViz) reveals close linkage between *baf* and *api* orthologs in many Gamma and Beta proteobacterial species, suggesting a functional link between them (60). This is consistent with our data indicating that Api and Baf might work together to inhibit LmdC auto processing (Fig. 7). One possibility is that Baf is the location-specific anchor for both Api and LmdC, and a coiled coil interaction between the periplasmic alpha helices of Api and LmdC prevents the catalytic CTD of LmdC from accessing the auto processing site (Fig. 8). LmdC auto processing can occur *in trans* because plasmid encoded LmdC-H282A, which is catalytically inactive, is processed in a *lmdC*^+^ strain (A.G. Sommerfield and A.J. Darwin, unpublished data). Nevertheless, a coiled coil Api-LmdC interaction could constrain the LmdC catalytic domain to prevent *in trans* auto processing. In addition to inhibiting auto processing, our experiments in *lmdC*-GG→AA mutant strains suggest that Api might also facilitate the function of unprocessed LmdC (Fig. 6b). AlphaFold predicts that the CTD of Api adopts a β-sandwich fold, a very common fold that often interacts with ligands, including proteins and other molecules. The CATH Protein Structure Classification Database places the Api β-sandwich domain in sub family 2.60.40.2420 of the Immunoglobulin-like superfamily, which includes CsgC/H-like proteins involved in formation of curli amyloid fibers. A Foldseek search using the AlphaFold Api structure as the query identifies several high probability matches to β-sandwich folds of proteins in the Protein data Bank. The top hits include CsgC/H proteins; FlaF from the Archaeal species *Sulfolobus acidocaldarius*; ApaG proteins, and the β-sandwich domain of *P. aeruginosa* BglX. These proteins have various functions that involve interactions: CsgC/H inhibit premature amyloid formation of curli fibers by interacting with the major fiber component CsgA; FlaF binds to FlaG and SlaA proteins to anchor the type IV pilus-like archaellum motility apparatus to the S-layer; ApaG proteins are thought to act as scaffolds for protein-protein interactions; the β-sandwich domain of BglX act as a structural scaffold and supports carbohydrate binding (61–64). The β-sandwich domains of ApaG and BglX form fibronectin type III (fn3)-like folds (63, 64). This fold has been found to interact with proteins, but also with sugar chains, including those of peptidoglycan (63, 65, 66). For all these reasons, our model proposes that the Api C-terminal β-sandwich domain might promote LmdC activity by interacting with LmdC itself and/or by anchoring to the cell wall target of LmdC (Fig. 8a)

*lmdC* and *baf* are not present in all *Pseudomonas* species, but when they are present one never occurs without the other, even though they appear in two different genomic contexts (Fig.6a). Absence from some *Pseudomonas* species, and different arrangements in others, might indicate separate acquisition events. *P. aeruginosa* LmdC is 63-64% identical to LmdC from *P. mendocina* and *P. fulva*, in which the *lmdC* genes are in the same context, but only 35-37% identical to LmdC from *P. stuzeri* (also now known as *Stutzerimonas stuzeri*) and *P. denitrificans* (also now known as *P.* sp. ATCC 13867), where *lmdC* occurs in a different genomic context. Similarly, *P. aeruginosa* Baf is 63-68% identical to Baf from *P. mendocina* and *P. fulva*, but only 33-35% identical to Baf from *P. stuzeri* and *P. denitrificans*. In *P. aeruginosa*, *P. mendocina* and *P. fulva* the gene immediately upstream of *baf* is *api* (Fig. 6a). However, the gene upstream of *baf* in *P. stuzeri* and *P. denitrificans* encodes a protein that appears to be a truncated form of Api. It is predicted to have the short cytoplasmic domain, transmembrane helix and periplasmic alpha helix found in Api, but it lacks the C-terminal β-sandwich domain. Representatives of this protein and Api both occur within the same two orthologous groups, ENOG502Z8KI and ENOG5032TUN, suggesting the possibility of divergence from a common ancestry. We speculate that either the role of the Api β-sandwich domain is not required in *P. stuzeri* and *P. denitrificans* or it is achieved by another unknown protein.

In summary, we have identified regulated LmdC auto processing preceding proteolytic destruction as a new mechanism of autolysin control. Amino acid sequences and predicted structures also suggest that LmdC auto processing is likely to be conserved beyond *P. aeruginosa*. We cannot yet make any conclusions about how Baf and Api inhibit auto processing and promote the function of LmdC, but the most likely explanations involve interactions. A predicted β-hairpin cytoplasmic domain is widely conserved in LmdC proteins, and this isolated part of *H. neptunium* LmdC interacted with bactofilin in an *in vitro* bio-layer interferometry assay (31, 39). Api also has a conserved small cytoplasmic tail that might interact with Baf. Detecting interactions between full length proteins *in vivo* would provide the most convincing evidence, but our attempts to detect LmdC/Baf/Api interactions with epitope-tagged pull downs and two-hybrid assays have so far yielded negative results. The majority of LmdC in the bacterial cell appears to be auto processed and degraded, suggesting that any association with Baf and Api involves a small minority of synthesized LmdC and that interactions could be weak and/or transient. Indeed, in some cases only weak two-hybrid interactions were observed and/or it has been difficult to detect robust bactofilin-partner interactions without having to use crosslinking followed by mass spectrometry for detection (51, 53, 67). Interaction of the *H. pylori* bactofilin CcmA with the Csd7 transmembrane protein could only be detected after cross-linking prior to co-immunoprecipitation and if mass spectrometry was used to detect CcmA, suggesting weak or transient Csd7-CcmA interactions (57). However, removal of the N-terminal region of CcmA led to robust co-purification with Csd7, suggesting that this CcmA region inhibits it from being co-immunoprecipitated by Csd7 (57). All of this suggests that a detailed, systematic investigation into LmdC/Baf/Api whole protein and domain interactions will be needed to rule this possibility in or out. This will be one of our future goals, along with investigating the locations of these proteins in individual cells and attempting to determine the precise effect of LmdC on peptidoglycan and of course its physiological role.

## MATERIAL AND METHODS

### Bacterial strains and growth conditions

Strains and plasmids used in this study are listed in Table 1. Routine growth was in Luria-Bertani (LB) broth, composed of 1% (wt/vol) tryptone, 0.5% (wt/vol) yeast extract, and 1% (wt/vol) NaCl or on LB agar. To select *P. aeruginosa* exconjugants after plasmid conjugation from *E. coli* strain SM10 donor, bacteria were recovered on Vogel-Bonner minimal agar with appropriate antibiotics (68, 69). Routine antibiotic concentrations were 200 μg/ml ampicillin for *E. coli*, 150 μg/ml carbenicillin for *P. aeruginosa,* 15 μg/ml tetracycline or gentamicin for *E. coli*, and 75 μg/ml tetracycline or gentamicin for *P. aeruginosa*.

**TABLE 1.**
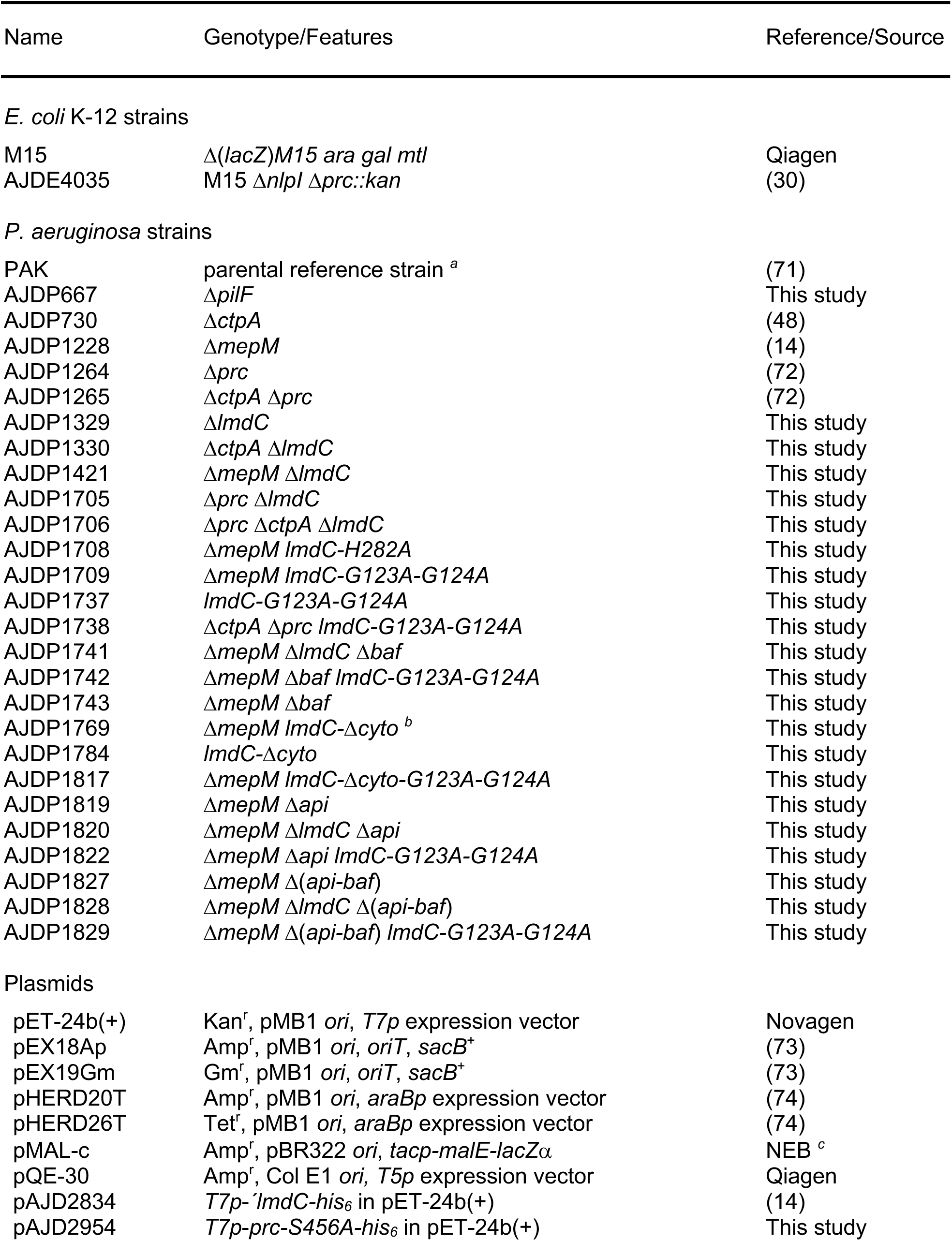
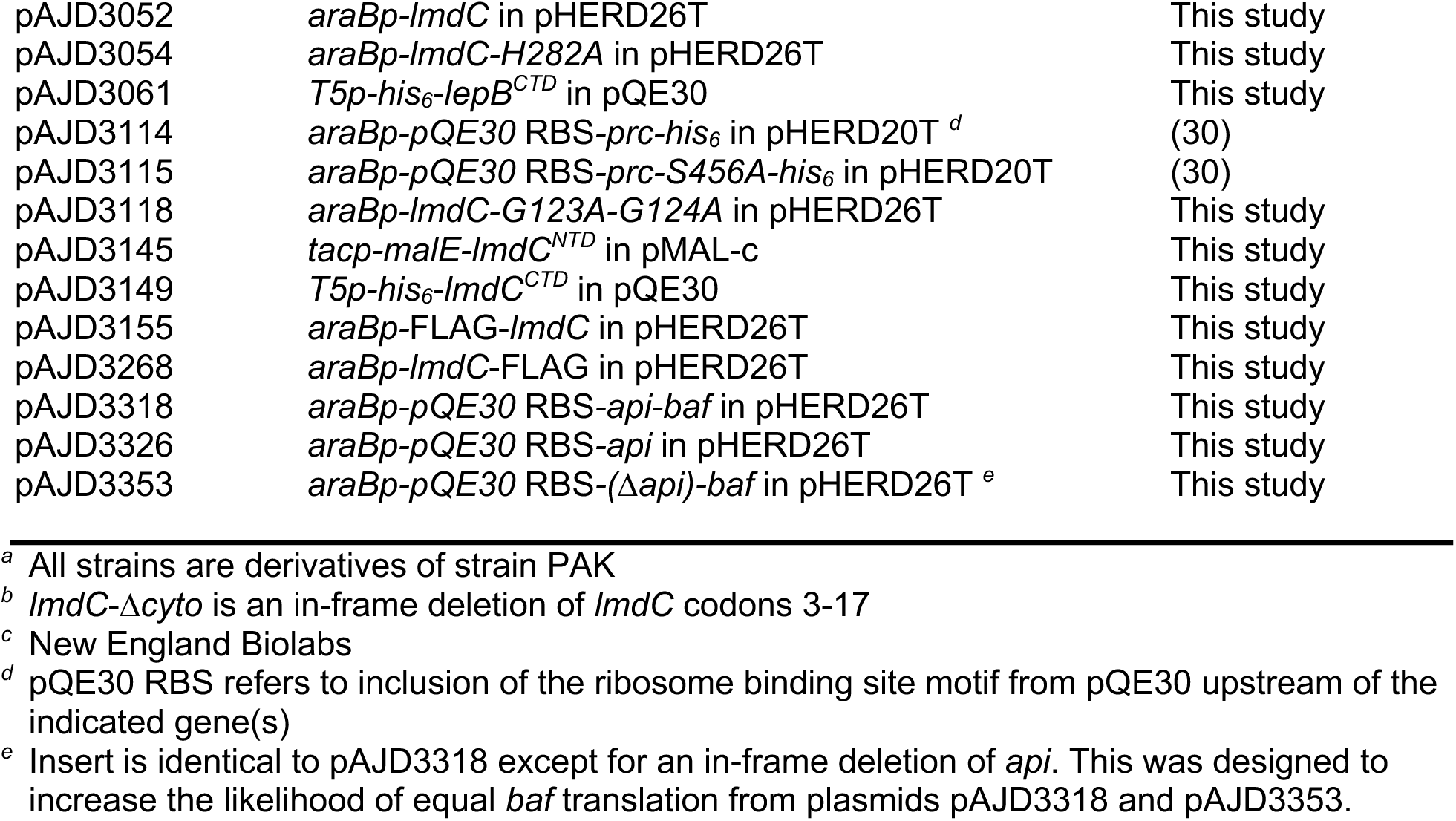
Strains and plasmids.

### Plasmid and strain constructions

To construct Δ*lmdC*, *lmdC*-Δ*cyto*, Δ*baf*, Δ*api*, Δ(*api-baf*) in frame deletion mutations, fragments of ∼0.55 kb corresponding to the regions upstream and downstream of the deletion site were amplified by PCR and cloned into pEX18Ap or pEX19Gm. These plasmids were integrated into the *P. aeruginosa* chromosome after conjugation from *E. coli* SM10 and sucrose-resistant carbenicillin- or gentamicin-sensitive segregants were isolated on LB containing 10% sucrose without NaCl. Deletions were verified by PCR analysis of genomic DNA.

For strains with *lmdC* G123A-G124A or H282A mutations, two ∼ 0.55 kb fragments flanking codons 123-124 or 282 of *lmdC* were amplified by PCR. For each fragment one of the primers incorporated mismatches at the target codon(s) to convert them to encode alanine. The two fragments were joined in a PCR SOEing reaction (70) via their overlapping regions around codons 123-124 or 282. The product was cloned into pEX18Ap and exchanged for the corresponding region of the *lmdC* gene by integration, selection for sucrose-resistant segregants and confirmation by colony PCR and DNA-sequencing.

pHERD26T expression plasmids were constructed by amplifying genes from *P. aeruginosa* chromosomal DNA using one primer that annealed upstream of the ribosome binding site and a second primer that annealed immediately downstream of the stop codon. However, for the *api*, *api-baf,* Δ(*api*)-*baf* expression plasmids the forward primer annealed immediately upstream of the *api* start codon and incorporated the strong ribosome binding site motif from plasmid pQE-30. This was done because *api* is translationally coupled to the upstream *argC* gene in the PAK genome and does not have its own obvious ribosome binding site. For the lmdC-FLAG plasmid the reverse primer incorporated a FLAG tag-encoding sequence immediately upstream of the stop codon. For the FLAG-*lmdC* plasmid the forward primer incorporated a FLAG tag-encoding sequence immediately downstream of the start codon. All amplified fragments were cloned into pHERD26T using restriction sites incorporated by the PCR primers.

For LmdC^NTD^ and LmdC^CTD^ proteins for *in vitro* proteolysis reactions, a plasmid encoding MBP-LmdC^NTD^ was made by amplifying codons 47-123 of *lmdC* by PCR. The upstream primer incorporated an EcoRV site for in-frame fusion to the *malE* gene of plasmid pMAL-c. The downstream primer incorporated a stop codon followed by an XbaI site. The PCR fragments were digested with EcoRV/XbaI and ligated into plasmid pMAL-c plasmid that was digested with StuI/XbaI. The plasmid encoding His_6_-LmdC^CTD^ was made by amplifying a fragment encompassing codon 124 to the stop codon by PCR and cloning it into pQE-30 as a BamHI/SacI using restriction sites incorporated by the PCR primers.

To produce antigens for polyclonal antiserum production, a plasmid encoding Prc-S456A-His_6_ was made by amplifying *prc-S456A* without the region encoding its signal sequence by PCR and cloning it into pET-24b(+) as a BamHI/XhoI fragment using restriction sites incorporated by the PCR primers. A plasmid encoding His_6_-LepB^CTD^ was made by amplifying codons 84-stop of *lepB* by PCR and cloning it into pQE-30 as a BamHI/HindIII fragment using restriction sites incorporated by the PCR primers.

All plasmids constructed using PCR amplification were checked by DNA sequencing of the entire PCR-generated insert to ensure that no spurious mutations had been introduced.

### Polyclonal antisera production and immunoblotting

*E.coli* strain ER2566 (NEB) containing plasmid pAJD2834 encoding LmdC-His_6_ or pAJD2954 encoding Prc-S456A-His_6,_ or strain M15 [pREP4] (Qiagen) containing plasmid pAJD3061 encoding His_6_-LepB, were grown in LB broth to mid-log phase at 37°C with aeration. Protein production was induced with 1 mM IPTG for 3-4 h at 37°C. The proteins were purified under denaturing conditions by nickel-nitrilotriacetic acid (NTA)-agarose affinity chromatography as described by the manufacturer (Qiagen). Polyclonal rabbit antisera were raised by Labcorp Early Development Laboratories Inc. Production of the OprF antiserum was described previously (14).

Samples were separated by SDS-PAGE and transferred to nitrocellulose by semi dry electroblotting. Chemiluminescent detection followed incubation with a LmdC, Prc, LepB or OprF polyclonal antiserum or anti-FLAG (Sigma) or -MBP monoclonal antibody (NEB), then goat anti-rabbit IgG (Sigma) or goat anti-mouse IgG (Sigma) horseradish peroxidase conjugates used at the manufacturers recommended dilution. In some cases, band intensities were quantified using ImageJ 1.46r software (National Institutes of Health, Bethesda, Maryland, USA).

### Anti-FLAG immunoprecipitation

Bacteria were inoculated into 100 ml of LB broth with 75µg/mL tetracycline in a 250-ml flask to an OD_600_ of 0.05 and grown at 37°C with aeration for 5 h. Equivalent amounts of bacterial cells from all strains were collected by centrifugation, washed in 10 mM Tris-HCl, 10% (wt/vol) glycerol (pH 7.5), and resuspended in 2 ml nondenaturing lysis buffer (NDLB) (50 mM Tris-HCl, 300 mM NaCl, 5 mM EDTA, 10% [wt/vol] glycerol [pH 7.5]). Roche complete protease inhibitors were added, cells were disrupted by sonication, and 1% (wt/vol) lauryldimethylamine *N*-oxide (LDAO) was then added, followed by incubation with rotation for 1 h at 4°C. Insoluble material was removed by centrifugation at 16,000 × *g* for 30 min at 4°C. Thirty microliters of anti-FLAG M2 affinity resin (Sigma-Aldrich) in NDLB was added to the supernatant, and the mixture was incubated for 2 h at 4°C with rotation. A 1-ml spin column (Pierce catalog number 69725) was used to wash the resin 10 times with 500 μL NDLB containing 0.1% (wt/vol) Triton X-100. Proteins were eluted by incubating the resin in 100 μL of 400 μg/mL 3× FLAG peptide in NDLB containing 0.1% (wt/vol) Triton X-10 and rotating at 4°C for 30 min.

### Mass spectrometry analysis of full LmdC-His_6_ proteins purified from *E. coli*

Excised protein bands were cut in ∼1 cubic mm and destained with 1:1 (v/v) methanol and 100 mM ammonium bicarbonate at 4°C with agitation. Destained gel pieces were dehydrated first with a 100% acetonitrile rinse and subsequently further dried in a SpeedVac concentrator. Proteins were then reduced with 20mM dithiothreitol (Sigma) for one hour at 57⁰C at pH 7.5. After samples cooled to room temperature, they were alkylated with 50 mM iodoacetamide (Sigma) for 45 minutes at room temperature in the dark. The gel pieces were then resuspended in 0.6 M Sodium cyanoborohydride in triethylammonium bicarbonate (TEAB) and double the volume of 4% formaldehyde solution was added and incubated for 1 hour. The reaction was quenched with 1% ammonia. The gel pieces were washed twice with TEAB and again dehydrated using 100% acetonitrile followed by further dehydration in a Speedvac concentrator. The dried gel pieces were rehydrated using 100 mM ammonium bicarbonate solution with 500 ng trypsin (Promega) for overnight digestion with light agitation. A solution of 5% formic acid and 0.2% trifluoroacetic acid (TFA) R2 50 μm Poros (Applied Biosystems) beads slurry in water was added to the gel pieces at a volume double of that used for the digestion. The gel pieces were put on a shaker at 4°C for 2 hours. The Poros beads were loaded onto equilibrated C18 ziptips (Millipore), with 0.1% TFA, using a microcentrifuge for 30 seconds at 6,000 rpm. The beads were washed first with 0.1 % TFA and subsequently three times with 0.5% acetic acid. Peptides were eluted with 40% acetonitrile in 0.5% acetic acid followed by 80% acetonitrile in 0.5% acetic acid. The organic solvent was removed using a SpeedVac concentrator. The samples were reconstituted in 0.5% acetic acid and stored at -80⁰C until analysis.

An aliquot was loaded onto an Acclaim PepMap trap column (2 cm x 75 µm) in line with an EASY-Spray analytical column (50 cm x 75 µm ID PepMap C18, 2 μm bead size) using the auto sampler of an EASY-nLC 1200 HPLC (Thermo Fisher Scientific) with solvent A consisting of 2% acetonitrile in 0.5% acetic acid and solvent B consisting of 80% acetonitrile in 0.5% acetic acid. Peptides were gradient eluted into a Thermo Fisher Scientific Q Exactive mass spectrometer using the following gradient: 5 - 35% in 60 min, 35 - 45% in 10 min, followed by 45 - 100% in 10 min. High resolution full MS spectra were recorded with a resolution of 70,000 at m/z 200, an AGC target of 1e6, with a maximum ion time of 120ms, and a scan range from 400 to 1500m/z. The top 20 MS/MS spectra were collected with a resolution of 17,500, an AGC target of 5e4, maximum ion time of 120ms, one microscan, 2 m/z isolation window, and Normalized Collision Energy (NCE) of 27 and a dynamic exclusion of 30 seconds. The MS/MS spectra were searched against the LmdC sequence using Byonic (Protein Metrics). The search parameters were: precursor mass tolerance ±20 ppm, fragment mass tolerance ±0.02 Da, digestion parameters nonspecific (no enzyme restriction), fixed modification of carbamidomethyl on cysteine, variable modification of oxidation on methionine, deamidation on glutamine and asparagine, dimethyl on N-termini, proline, lysine and arginine, acetylation on protein N-term, methylation on lysine and arginine, and trimethylation on lysine. Peptides were filtered using a Byonic score cut off >300. N-terminally modified peptides were verified by manual inspection of the MS/MS spectra.

### Protein purification and *in vitro* proteolysis assays

Prc-His_6_ and Prc(S456A)-His_6_ were purified previously (30). To purify MBP-LmdC^NTD^, *E. coli* strain M15 containing plasmid pAJD3145 was grown in 1 L of LB broth at 37°C with aeration until OD_600_ was between 0.6 and 1.0. Protein production was induced by adding 1 mM IPTG and incubating for 3 h at 37°C with aeration. Cells were resuspended in cold column buffer (20 mM Tris-HCl, pH 7.4, 200 mM NaCl, and 1 mM EDTA, pH 7.4). Roche complete protease inhibitor was added immediately as well as 1 mg/mL lysozyme. Cells were incubated on ice for 30 min and disrupted by sonication, and then, insoluble material was removed by centrifugation. The supernatant was added to a column containing 2.5 mL amylose resin (NEB). The resin was washed with 50 mL of column buffer, and proteins were eluted with 20 mL cold column buffer containing 10 mM maltose, in 1-mL fractions. Fractions 2 and 3 were combined and used for the assays without any further treatment.

His_6_-LmdC^CTD^ was purified differently because it was insoluble when overproduced in *E. coli*. *E. coli* strain M15 [pREP4] containing plasmid pAJD3149 was grown in LB broth to mid-log phase at 37°C with aeration. Protein production was induced with 1 mM IPTG for 3 h at 37°C. Bacterial cells were harvested by centrifugation and resuspended in 6 M guanidine hydrochloride, 100 mM NaH_2_PO_4_, 10 mM Tris-HCl, 5 mM imidazole, and 1% (wt/vol) Triton X-100 (pH 7.5) by stirring at room temperature for 1 h. The suspension was sonicated, and insoluble material was removed by centrifugation at 10,000 × *g* for 30 min. Ni-NTA-agarose was added to the supernatant and stirred for 1 h at room temperature. The Ni-NTA-agarose resin was collected in a drip column and washed with 30 ml of a solution 50 mM NaH_2_PO_4_, 300 mM NaCl, and 20 mM imidazole (pH 8) to renature the protein. Protein was eluted in 2 mL fractions using 50 mM NaH_2_PO_4_–300 mM NaCl (pH 8) buffer containing increasing imidazole concentrations (50, 100, 150, 200, and 250 mM). 200-250 mM imidazole fractions were combined and used in the assays without any further treatment.

*In vitro* proteolysis reactions contained approximately 2 µM Prc-His_6_ or Prc-S456A-His_6_ and approximately 2 µM MBP-LmdC^NTD^ or His-LmdC^CTD^. Reactions were incubated at 37°C, and aliquots were removed after 1, 2, 3, and 4 h and terminated by adding SDS-PAGE sample buffer and boiling. Proteins were separated by SDS-PAGE and stained with ProtoBlue Safe (National Diagnostics) or analyzed by anti-MBP or anti-LmdC immunoblot as described above.

### Subcellular fractionation

Bacteria were inoculated into 600 ml of LB broth in a 1 L flask to an OD_600_ of 0.05 and grown at 37°C with aeration until the OD_600_ was 0.6-0.8. Bacterial cells were collected by centrifugation and resuspended in 15 mL of cold 10mM Tris-HCL, 20% (w/w) sucrose pH 7.4. The suspension was supplemented with DNase I and RNase A, each to 10 µg/ml final concentration, as well as complete EDTA-free protease inhibitors (Thermo Scientific). Cells were lysed by passage through a high-pressure cell disruptor for 10 min and unbroken cells were removed by centrifugation at 2,000 x *g* for 15 min at 4°C. Membranes were collected from the supernatant by ultracentrifugation at 120,000 x *g* for 45 min at 4°C. The supernatant was retained as the soluble fraction and the membrane pellet was homogenized in approximately 5 ml of 20% (w/w) sucrose in 10 mM Tris-HCl, 1 mM EDTA pH 7.4. A small amount of the homogenate was retained as the total membrane fraction and the remainder was placed on top of a discontinuous sucrose gradient made by layering 1 mL each of 75%, 60%, 55%, 50%, 45%, 35 and 30% (w/w) sucrose in 10 mM Tris-HCl, 5 mM EDTA pH 7.4 into a 14 x 89 mm Ultra-Clear tube (Beckman Coulter). The gradients were centrifuged using a Beckman Coulter SW 41 Ti rotor at 274,000 x *g* for 48 h at 4°C. Twenty fractions (592 µl) were collected from the top of the tubes and analyzed by SDS-PAGE followed by immunoblotting.

### Growth Curves

Saturated cultures were diluted to OD_600_ of 0.01 in LB broth with or without ticarcillin-clavulanic acid in a total volume of 150 µL in the wells of a honeycomb plate. Plates were incubated in a Bioscreen C Automated Plate Reader (Oy Growth Curves Ab, Ltd.) at 37°C with continuous agitation and OD_600_ measurements every 15 min for 24 h.

### Twitching motility assay

Motility agar was made by pouring 25 mL of 1% (wt/vol) molten agar in 10% (wt/vol) tryptone, 5% (wt/vol) yeast extract and 5% (wt/vol) NaCl into petri plates and allowing them to solidify at room temperature for 1 h in a single layer. The plates were then inverted and left to dry overnight at room temperature. Test strains were streaked onto LB agar and grown at 37°C overnight. A small portion of the outer edge of the streak was removed with a plastic inoculation loop and mixed in a sterile area of the agar until it formed a smooth paste. A pipette tip was used to scoop up some of the paste, attempting to keep the inoculum size similar for each plate, and then stabbed perpendicular to the motility agar down to the agar-plastic interface in the center of the plate. The plates were inverted, placed into a humidified box in stacks of no more than three and incubated at 37°C for 40 h. The diameter of the halo of bacteria that had twitched across the plate between the bottom of the agar and the plastic was measured. A twitching-motility deficient Δ*pilF* mutant was assayed simultaneously as a negative control to ensure that the halo corresponding to twitching motility had been correctly identified.

## Data availability

The proteomic mass spectrometric raw files are accessible at https://massive.ucsd.edu under accession MassIVE , and at www.proteomexchange.org under accession -----. doi: , dataset license: CC0 1.0 Universal (CC0 1.0).

## ACKNOWLEDGEMENTS

Research and publication were supported by the National Institute of Allergy and Infectious Diseases (NIAID) of the National Institutes of Health (NIH), under award numbers R01AI136901, R21AI151097 and R01AI195551 to A.J.D., and award number T32AI007180 also supported A.G.S. The content is solely the responsibility of the authors and does not necessarily represent the official views of the NIH. The mass spectrometric experiments were done by NYU Grossman School of Medicine Proteomics Laboratory (RRID: SCR_017926) and partly supported by NYU Langone Health. We thank Dolon Chakraborty, Maria Chiara Mastropasqua, Terry-Ann Smith and Michelle Wang for constructing some of the strains and plasmids we used in this study, and Eliza Zaroff for comments on a draft version of the manuscript.

